# SWORD - a highly efficient protein database search

**DOI:** 10.1101/014654

**Authors:** Robert Vaser, Dario Pavlovi, Matija Korpar, Mile Sikic

## Abstract

**Motivation:** Protein database search is one of the fundamental problems in bioinformatics. For decades, it has been explored and solved using different exact and heuristic approaches. However, exponential growth of data in size in recent years has brought significant challenges in improving already existing algorithms. BLASTP has been the most successful tool for protein database search, but is also becoming a bottleneck in many applications. Due to that, many different approaches have been developed to complement or replace BLASTP. In this paper, we present SWORD, an efficient protein database search implementation that runs 3-4 faster than BLASTP in the sensitive mode and up to 18 faster in the fast and less accurate mode and also provides guaranteed optimal alignments for candidate sequences. SWORD is designed to be used in nearly all database search environments, but is especially suitable for large databases. Its sensitivity exceeds that of BLASTP for majority of input datasets.

**Availability:** Sword is freely available for download from https://github.com/rvaser/sword

## 1 INTRODUCTION

Protein database search is an immensely important task in bioinformatics and other life sciences. It is used both as a standalone task of finding similarities between proteins and, for example, as a part of larger metagenomical studies. However, a large and ever-growing amount of new data being analyzed together with an exponential increase of database sizes makes protein similarity analysis using existing tools a dauntingly slow and inefficient task. BLASTP (Altschul *et al.,* 1990) is the most popular and used software nowadays for protein database search with tens of thousands of searches on the NCBI servers a day (Liu *et al.,* 2011).

In short, BLASTP uses a so-called seed and extend approach in four stages. Firstly, BLASTP tries to find *hits*. Hits are short exact matches (usually of size 3) between query and target sequence that score above some small threshold according to the scoring matrix being used. In the second stage, these hits are extended using ungapped extension. All the extensions that score above a certain cutoff score are passed to the third and fourth stages. These are called high-scoring segment pairs (HSPs). In the third and fourth stage, alignment between the query and the target is computed using HSPs as seeds. All alignments that score above some predefined threshold are reported. Note that these alignments are not necessarily optimal since they are based on heuristic measures from the second stage. This process is rather time consuming on most large databases, such as NCBI nonredundant (NCBI-NR) that is 32GB in size.

For that reason, many new approaches and tools like DIAMOND (Buchfink *et al.,* 2014), BLAT (Kent, 2002) and PAUDA (Huson and Xie, 2014) have been developed in recent years with the goal of speeding up the search while retaining the same or at least a comparable level of sensitivity to that of BLAST. Many of these approaches try to make use of new and parallel computing architectures to help cope with huge computational demands while others present completely new ideas and perspectives on the problem.

One of them is DIAMOND, which is, according to its paper, up to 20 000 faster than BLAST in a high-throughput setting where an extremely large number of short reads needs to be aligned to a protein database. It is targeted as a replacement for BLASTX in such an environment. DIAMOND uses a few novel ideas in its approach, most notable being double indexing and spaced seeds. Double indexing helps in reducing memory load by being more cache conscious which in turn significantly improves the runtime, while spaced seeds were created with the intention of locating hits in a more efficient manner by having a need to process much less seeds than usual without losing sensitivity. Hits are then passed to SIMD-accelerated banded Smith-Waterman algorithm for alignment. However, being specifically designed to be extremely fast and accurate as much as possible when aligning a huge number of reads to a protein database, makes DIAMOND sensitivity somewhat poor on regular, longer query data sets, which we will examine and present later on.

Other tools, such as BLAT, Rapsearch2 (Zhao *et al.,* 2012) or PAUDA propose other, different approaches which can yield good results depending on the application, with sometimes very big speedups over BLASTP and purely exact algorithms. Nonetheless, efficient and sensitive protein database search still remains a challenge to be solved. We have used BLASTP and DIAMOND for comparison with SWORD.

In this paper, we present SWORD or Smith-Waterman On Reduced Database - a novel approach to this problem that combines some well-known ideas with some new insights and possibilities opened up by hardware architectures. SWORD is, depending on parameters used, up to 18 times faster than conventional BLASTP in the same setting while providing clients with comparable sensitivity. It makes use of a heuristic preprocessing phase for reducing the search space followed by the second, optimal alignment phase. The second phase is sped-up by Nvidia CUDA enabled GPUs that run full Smith-Waterman algorithm, thus providing user with optimal alignments.

## 2 METHODS

We have implemented a two-step approach to database searching that involves a heuristic and an optimal alignment phase. In the heuristic part, we use deterministic finite automata to eliminate from further processing those database sequences which probably will not produce good alignments and therefore we reduce the number of alignments computed in the second phase, thus saving time. In the second step, we use SW#db (Korpar *et al.,* 2015), an efficient Smith-Waterman algorithm library that uses CUDA GPU programming parallel architecture. SW#db makes use of computing power of CUDA cards paired with multithreading and SIMD vectorization to help make running larger exact alignments feasible.

While BLASTP (and the similar algorithms) use the results from the seed-and-extend phase to build final alignments, our approach only uses the heuristic seed phase to find possible target matches. Final alignments are independently calculated using SW and are guaranteed to be optimal.

The heuristic we use for determining possible matches is based on a deterministic finite automaton (DFA). The DFA is built to recognize equal-length k-mers extracted from the query sequences, called seeds. The seeds need not be just the exact kmers from the query sequence, but can also include ‘similar enough’ k-mers. The similarity between the seeds is defined by the threshold and the scoring matrix used. The DFA is used to locate *hits:* occurrences of query seeds in the target sequence. For a pair of k-mers, one from the query and one from the target, to be considered a hit, they have to score at least *T*, where *T* is a predefined threshold. All seeds that score *T* or more with a given seed belong to its *neighborhood.* Detecting not only equal, but all seeds from a seed neighborhood greatly increases sensitivity. SWORD supports both modes of operation. To enable detection of similar, neighborhood seeds, we use the following technique. While iterating through query seeds and adding them to the DFA, for each seed we generate all seeds that score *T* or higher when aligned to it, and insert them to the automaton, too. This is very similar to how BLASTP used to detect HSPs.

When a hit is found using the DFA, we calculate the *diagonal* on which the hit is located. If a query is of length *M* and a target is of length N, then a diagonal is defined as a diagonal of an *MxN* matrix. The following formula determines the diagonal on which a hit is located:

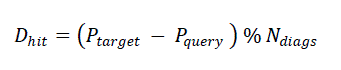

where *D_hit_* is the index of the diagonal for the current hit, *P_target_* and *P_query_* are hit positions in the target and the query respectively while *N_diags_* represents total number of different possible diagonals between the current query and the current target and is obtained by the following expression:

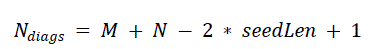

Here, *seedLen* is the length of seeds used while constructing the automaton. When the diagonal is calculated, we increment the count of the number of hits on that diagonal. In the end, after processing the current target is done, the diagonal with the largest number of hits is found and used as the relevance score or probability of that target being potentially good. We use this measure because more hits on the same diagonal indicate an increase in the probability of a good alignment between sequences.

The result of the heuristic is, for each target, a measure of probability of producing a good alignment with the query. Targets are then sorted by that measure in descending order and only the top *A* are sent to the second step for further analysis and processing, *A* being an input parameter of SWORD. Those *A* targets are now called *candidates* since all of them will be aligned to the query, but only a predefined number of them will be reported. In the second step, an exact Smith-Waterman alignment is performed between the query and all the candidate sequences, after which the alignments are filtered by score and expect value and reported in the output.

Counting hits on diagonals is method used in original FASTA (Pearson and Lipman, 1988) algorithm. However, FASTA combines it with the other processing steps before moving on to alignment/extend phase. We, on the other hand, use only hit count as a relevant measure. This produces more targets to align against and does not reduce the search space as much as FASTA does, but we rely on the power and efficiency of SW#db and CUDA cards to compensate for the expansion of the search space.

We have implemented additional improvements to the described algorithm to make the program even more efficient. Firstly, we group multiple queries into one automaton. In the default implementation, it is necessary to iterate over the whole database for each query in order to detect hits and find the best possible candidates. If multiple queries are grouped into one automaton, we can detect hits for all of those queries in a single iteration. We were careful to implement the grouping in a cache friendly manner. The software is also optimized for multi-core processors.?

SWORD supports two modes of operation – fast and sensitive. As the names suggest, the former is faster and provides lower sensitivity rates, while the latter gives larger sensitivity at a cost of less speedup. Users can manually set up their own preferred way of database search with many other parameters available.^1^ The sensitive mode of operation uses seeds of length 3 along with generating and detecting neighborhoods with threshold of *T* = 13. It will forward the 30000 candidates per query to the second stage. On the other hand, fast mode uses seeds of length 5 and detects only exact matches. Only 5000 best targets per query are forwarded to the second stage as candidates in the fast mode.

It is important to note that SWORD needs no additional preprocessing of queries or the database before running. BLASTP and DIAMOND, for example, need to create their own auxiliary database files. SWORD requires no such files and takes up no extra disk space.

## RESULTS

We have tested and compared SWORD's performance to both BLASTP and DIAMOND. We have downloaded the NCBI NR and used it as the reference database. In order to better evaluate and compare sensitivity and speed, we have used three different query sets, two of them for sensitivity checks and all three to test runtime. The three sets were HumDiv, HumVar (Adzhubei *et al.,* 2010), both neutral variants and the third was E.coli (Flicek *et al.,* 2013) from Ensembl gene annotation. HumDiv contains 315 proteins of total length of 235219 amino-acids. HumVar is made of 3400 sequences of total length 2263906 acids while E.coli set contains 4969 protein sequences of total length 1393750.

Since SWORD and DIAMOND support fast and sensitive modes, we executed both variants for them. BLOSUM62 was the scoring matrix used in all runs. We ran BLASTP with default parameters, except that we turned off composition based statistics. Since we don't change the scoring matrix during or just prior to the search and don't use composition based statistics, turning them off makes BLASTP behave in the same manner. This enabled us to get more comparable e-values for alignments since they are all now calculated in the same way. All programs were run using 32 threads on a machine with 16 Intel(R) Xeon(R) E5-2640 CPUs clocked at 2GHz with HyperThreading support, 256 GB SSD and 396GB RAM. GPU unit used for testing was Nvidia Geforce GTX Titan. The BLASTP and DIAMOND auxiliary files for NR database, which is around 32GB in size, occupied about 60GB of extra disk space and took an extra few hours to be computed.

To measure sensitivity we used HumDiv and E.Coli sets. First, we ran SW#db search and stored the alignments produced. We proceeded to run SWORD, DIAMOND and BLASTP on the same sets. Sensitivity was measured by examining 11 different e-values, from 1*x*10^−250^ to 10. For each of those threshold e-values and for each program we found all alignments that have e-value less than or equal to the threshold. We then examined what percentage of those produced alignments is present in the SW#db exact output and reported the percentage as sensitivity for the given e-value. The results of these tests can be seen on Figure for both input sets.

Horizontal axis represents e-values while the vertical axis represents the percentage of SW#db alignments covered by an algorithm. For every algorithm and e-value, this percentage is plotted. From these figures, we can easily observe that SWORD in its sensitive variant has very good overall sensitivity. It does perform slightly worse than BLASTP on higher e-values for E.coli, but that is probably due to a larger number of mutations occurring in bacteria. These mutations are harder to detect, especially since we use larger neighborhood threshold than BLASTP (13 compared to 11 in BLASTP) Also, SWORD-fast performs quite well, although its sensitivity drops more after a certain point. The reason for this is that we are only looking at exact matches of length 5 without checking seed neighborhoods. On very similar sequences with low e-value alignments this works well since they probably share at least one exact seed of size 5. However, when sequences are more distant, in many cases they share no exact seed matches of that size and SWORD-fast does not discover any hits between them. DIAMOND, on the other hand, performs somewhat poorly compared to SWORD and BLASTP, which is expected since it was designed for different scenarios. Figures also demonstrate that SWORD, in both variants, can be used to efficiently retrieve very similar proteins from the database with equal or better sensitivity than BLASTP but with speedup of 3 or more times. This usually is the primary task when searching a database.

During execution speed testing, we tested how much speedup DIAMOND and SWORD provide compared to BLAST. These results can be seen on Figure 2. A few things can be read from the graph. As expected, when the query set increases in the number of queries and/or their size, the speedup of SWORD over BLASTP becomes larger. This means that it scales well with the data set size and is well suitable for large sets. SWORD was able to surpass both DIAMOND counterparts on the HumDiv set. DIAMOND is very fast on all sets, but we are not sure how it scales. Its performance seems to dip significantly on HumDiv set and it is hard to conclude why.

**Figure 1.**
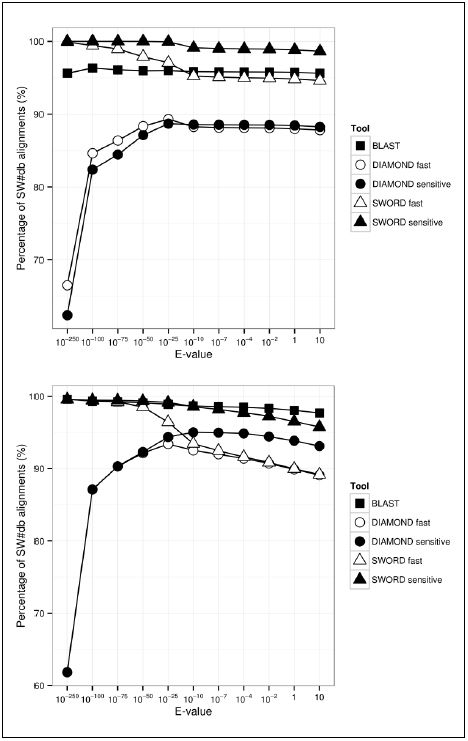
Comparison of sensitivity of SWORD against BLASTP and DIAMOND on HumDiv (upper) and E.coli set (lower) for NR database.

**Figure 2.**
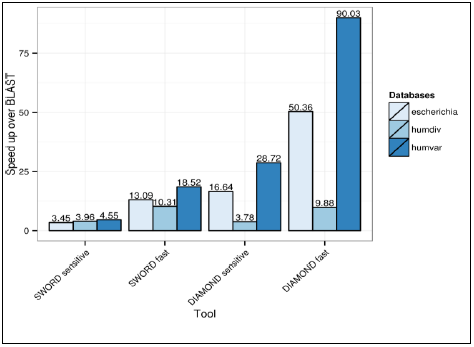
Comparison of speedup over BLAST for all algorithms on three sets for NR database.

In addition, we have compared SWORD to DIAMOND and BLASTP with SW#db as reference on the Astral/SCOP compendium database, version 2.04 (Fox *et al.,* 2014). For this testing, we created a query set from the subset of Astral sequences. The query set was created by sorting the SCOP domains in lexicographic order and selecting even numbered sequences as queries. The database consisted of 13042 sequences while the query set contained 6114 sequences. Curves denoting the number of true positives vs. the number of false positives for each algorithm are plotted on Figure 3. DIAMOND results are poor and it finds only a fraction of all alignments. SWORD-sensitive performance is drawn using red dashed line that almost covers the curve of SW#db completely. From this, it is obvious that SWORD-sensitive performs almost exactly like SW#db in this case, while other algorithms are significantly less accurate.

**Figure 3.**
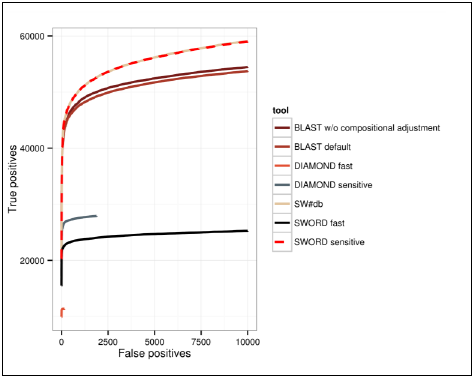
Comparison of sensitivity of SWORD, BLASTP and SW#db on Astral/SCOP database.

## ACKNOWLEDGEMENTS

We would like to give special thanks to Ana Bulovic for proofreading the manuscript and providing valuable suggestions.

*Funding:* This work was supported in part by Croatian Science Foundation under the project 7353 Algorithms for Genome Sequence Analysis.

## Footnotes

1 A detailed description of SWORD parameters is available on the download page

